# SF3B1-dependent alternative splicing is critical for maintaining endometrial homeostasis and the establishment of pregnancy

**DOI:** 10.1101/2023.05.20.541590

**Authors:** Pooja Popli, Sangappa B. Chadchan, Michelle Dias, Xinchao Deng, Stephanie J. Gunderson, Patricia Jimenez, Hari Yalamanchili, Ramakrishna Kommagani

**Affiliations:** Department of Pathology and Immunology; Department of Pediatrics, Neurology, Baylor College of Medicine, One Baylor Plaza, Houston, TX, 77030, USA; Center for Reproductive Health Sciences, Washington University School of Medicine, St Louis, MO, USA; Department of Obstetrics and Gynecology, Washington University School of Medicine, St Louis, MO, 63110, USA; Department of Molecular Virology and Microbiology, Baylor College of Medicine, One Baylor Plaza, Houston, TX 77030, USA

**Keywords:** Endometrium, embryo implantation, splicing factor, pladienolide-B, alternative splicing, mRNA maturation.

## Abstract

The remarkable potential of human endometrium to undergo spontaneous remodeling is shaped by controlled spatiotemporal gene expression patterns. Although hormone-driven transcription shown to govern these patterns, the post-transcriptional processing of these mRNA transcripts, including the mRNA splicing in the endometrium is not studied yet. Here, we report that the splicing factor, SF3B1 is central in driving alternative splicing (AS) events that are vital for physiological responses of the endometrium. We show that loss of SF3B1 splicing activity impairs stromal cell decidualization as well as embryo implantation. Transcriptomic analysis revealed that SF3B1 depletion decidualizing stromal cells led to differential mRNA splicing. Specifically, a significant upregulation in mutually exclusive AS events (MXEs) with SF3B1 loss resulted in the generation of aberrant transcripts. Further, we found that some of these candidate genes phenocopy SF3B1 function in decidualization. Importantly, we identify progesterone as a potential upstream regulator of SF3B1-mediated functions in endometrium possibly via maintaining its persistently high levels, in coordination with deubiquitinating enzymes. Collectively, our data suggest that SF3B1-driven alternative splicing plays a critical role in mediating the endometrial-specific transcriptional paradigms. Thus, the identification of novel mRNA variants associated with successful pregnancy establishment may help to develop new strategies to diagnose or prevent early pregnancy loss.

## Introduction

Alternative splicing is a fundamental process by which a single gene generates multiple distinct mRNA transcripts and thereby increases protein diversity. Over 90% of human genes undergo alternative pre-mRNA splicing catalyzed by the spliceosome (1–3), a macromolecular complex composed of five small nuclear RNAs (U1, U2, U4, U5, U6) and more than 200 additional proteins (4–6). The first step of mRNA splicing is the recognition of specific sequences on pre-mRNA by spliceosomal proteins. Next, introns and various combinations of exons are removed to generate mature mRNA variants containing different exon combinations (7). Thus, failure to accurately recognize splice sites due to splice site mutation or splicing factor dysregulation leads to the generation of abnormal mature mRNA variants (8, 9) and cellular dysfunction (10–22). These aberrantly generated mRNA variants can act as drivers of several human diseases, including cardiovascular disease, Alzheimer’s disease, and cancers (9, 23–26).

SF3B1 is among the principal regulators of alternative splicing and functions as a core component of the U2 snRNP spliceosome complex, which is critical for branch site recognition and for the early stages of spliceosome assembly (5). Missense mutations in this gene are frequently found in several different blood cancers (e.g., myelodysplastic syndrome, chronic lymphocytic leukemia) and solid tumors (e.g., breast and pancreatic cancers, uveal melanoma) (27). Recent studies including ours have also found a link between wild-type SF3B1 expression levels and tumor aggressiveness in pancreatic, endometrial, prostate, liver, and breast cancers, with higher expression being associated with poor prognosis (9, 28–31). Importantly, SF3B1 germline deletion results in perinatal or embryonic lethality, indicating a role for SF3B1 in the developmental process (32). Despite the indispensable role of SF3B1 during early development, its functions in adult tissues, including the endometrium, are largely unknown. Uncovering the precise roles of SF3B1 in human physiologies is thus an important question to investigate.

Up to 37% of conceptions fail to advance beyond 20 weeks of gestation. Additionally, up to 5% of women experience recurrent pregnancy loss, defined as two or more failed clinical pregnancies. Although some cases of early pregnancy loss are due to embryo abnormalities, up to 60% of cases involve normal blastocysts, indicating that those pregnancy losses are due to impaired uterine function leading to failed embryo implantation. To prepare for implantation, the uterus undergoes several well-timed estrogen and progesterone-mediated cellular changes (33, 34). Much of our understanding on these cellular changes are gleaned from rodent studies (35). In early pregnancy mice, estrogen promotes uterine epithelial proliferation, then, progesterone causes epithelial proliferation to stop, and the uterus become conducive for embryo attachment. In response to embryo attachment to uterine epithelium, the underlying stromal cells differentiate into decidual cells (decidualization), which determines the depth of trophoblast invasion and placentation (36–38). A defect in any of these processes can lead to implantation failure and early pregnancy loss (39). Thus, to develop strategies to diagnose and prevent early pregnancy loss, we must fully define the molecular mechanisms underlying these hormone-driven processes.

In this study, we present data that underscores the importance of progesterone-mediated SF3B1-driven alternative splicing in endometrial homeostasis. We show that SF3B1 protein levels are modulated during the early stages of pregnancy, and pharmacological inhibition of SF3B1 activity results in impaired stromal cell decidualization. Furthermore, we report that loss of SF3B1 causes robust alteration in splicing events, resulting in generation of various aberrant transcript variants in stromal cells and eventually hampering decidualization. Finally, we define the molecular mechanism for how progesterone acts as an upstream regulator and stabilizes SF3B1 levels in stromal cells during the process of decidualization.

## Materials and Methods

### Animal care and use

All animal studies were approved by the Institutional Animal Care and Use Committees (IACUC) of Washington University School of Medicine, Saint Louis, MO, USA and Baylor College of Medicine, Houston, TX, USA. C57BL/6 Taconic mice from Jackson Laboratories were maintained in standard 12-h light/dark conditions and provided ad libitum access to food and water. Animals were handled according to an approved IACUC protocol number [AN-8890].

### Artificial induction of decidualization

Decidualization was artificially induced using previously described methods (40–42). In brief, six-week-old female WT mice were bilaterally ovariectomized under ketamine anesthesia and buprenorphine-SR as analgesics. Mice were thereafter rested for two weeks to allow endogenous ovarian hormones to dissipate. Mice were subcutaneously injected with 100 ng estrogen (E2) on the next three consecutive days. After two days of rest, the mice were subcutaneously injected with 1 mg progesterone (P4) and 10 ng E2 on each of the next five days. Six hours after the third E2+P4 injection, 50 µL of sesame oil was injected into the lumen of the right uterine horn to induce decidualization. The other uterine horn was left untreated as a control. After oil instillation, mice received daily injections of 10 ng E2 plus 1 mg P4 (E2+P4) along with vehicle of PLAD-B on each of the next two days while the E2+P4 treatment continued for one more day. The mice were euthanized six hours after the last injection, and the wet weights of the left and right uterine horns of each mouse were recorded. Both uterine horns were fixed in 4% neutral buffered paraformaldehyde, snap frozen, and stored at −80°C.

### Human endometrial stromal cell isolation

For human endometrial biopsies, informed consent was obtained in accordance with a protocol approved by the Washington University in St. Louis Institutional Review Board (IRB ID #: 201612127). Endometrial biopsies from healthy women of reproductive age were obtained during the proliferative phase (days 9 to 12) of the menstrual cycle (43, 44). Human endometrial stromal cells (HESCs) were isolated as described previously (44). All experiments with human endometrial stromal cells were done independently in three replicates of cells from three patients.

### Transfection and HESCs decidualization

HESCs were seeded in six well culture plates at a cell density of 1×10^5^ per well in triplicates. For siRNA-mediated knockdown of *SF3B1*, HESCs were treated with Lipofectamine RNAiMAX reagent and 60 pmol of either non-targeting siRNA (D-001810-10-05) or siRNAs targeting *SF3B1* (L-020061-01-0005). Six hours post-transfection, media was replaced with HESC media. Forty-eight hours later, HESCs were treated with Opti-MEM reduced serum media containing 2% fetal bovine serum (FBS), E2 (100 nM), medroxyprogesterone acetate (MPA: 10 mM (Sigma-Aldrich) and cAMP (50 mM (Sigma-Aldrich)) which constitutes the decidualization media. The first day that HESCs were cultured in decidualization media was assigned day 0. Decidualization media was renewed every two days. Cells were harvested at appropriate time points as per experimental conditions. Total RNA was isolated to assess transcript levels of the decidualization markers: prolactin (PRL) and insulin-like growth factor binding protein-1 (IGFBP-1) (41).

### Timed mating and visualization of implantation sites

Sexually mature mice (6 weeks of age or older) were mated to fertile wild-type males and copulation was confirmed by the observation of vaginal plugs the following morning. The morning when the plug was observed is designated as 1 dpc. Implantation sites were visualized in 5 dpc pregnant mice by tail-vein injection of 50 µL of 1% Chicago Blue B. Dissected uteri were photographed, and the blue beads indicating implantation sites were counted.

### Histological analysis

The collected uterine tissues were fixed in 4% paraformaldehyde and embedded in paraffin. Sections (5 μm) were immunostained (n = 5 per group) (45). Briefly, after deparaffinization, sections were rehydrated in an ethanol gradient, then boiled for 20 min in citrate-buffer (Vector Laboratories Inc., CA, USA) for antigen retrieval. Endogenous peroxidase activity was quenched with Bloxall (Vector Laboratories Inc., CA, USA), and tissues were blocked with 2.5% goat serum in PBS for 1 hr (Vector Laboratories Inc., CA, USA). After washing in PBS three times, tissue sections were incubated overnight at 4°C in 2.5% goat serum containing the primary antibodies listed in **Table S1**. Sections were incubated for 1 hr with biotinylated secondary antibody, washed, and incubated for 45 min with ABC reagent (Vector Laboratories Inc., CA, USA). Color was developed with 3, 3′-diaminobenzidine (DAB) peroxidase substrate (Vector Laboratories Inc.), and sections were counter-stained with hematoxylin. Finally, sections were dehydrated and mounted in Permount histological mounting medium (Fisher Scientific).

### Immunofluorescence

Formalin-fixed and paraffin-embedded sections from both proliferative or secretory endometrium biopsies (n=5) were deparaffinized in xylene, rehydrated in an ethanol gradient, and boiled in citrate-buffer (Vector Laboratories Inc., CA, USA) for antigen retrieval. After blocking with 2.5% goat-serum in PBS (Vector laboratories) for 1 hr at room temperature, sections were incubated overnight at 4°C with primary antibodies (**Table S1**) diluted in 2.5% normal goat serum. After washing with PBS, sections were incubated with Alexa Fluor 488-conjugated secondary antibodies (Life Technologies) for 1 hr at room temperature, washed, and mounted with ProLong Gold Antifade Mountant with DAPI (Thermo Scientific).

### Western blotting

Protein lysates (40 μg per lane) were loaded on a 4-15% SDS-PAGE gel (Bio-Rad), separated in 1X Tris-Glycine Buffer (Bio-Rad), and transferred to PVDF membranes via a wet electro-blotting system (Bio-Rad), all according to the manufacturer’s directions. PVDF membranes were blocked for 1 hr in 5% non-fat milk in Tris-buffered saline containing 0.1% Tween-20 (TBS-T, Bio-Rad), then incubated overnight at 4°C with antibodies listed in **Table S1** in 5% BSA in TBS-T. Blots were then probed with anti-Rabbit IgG conjugated with horseradish peroxidase (1:5000, Cell Signaling Technology) in 5% BSA in TBS-T for 1 hr at room temperature. Signal was detected with the Pierce ECL Western Blotting Substrate (Millipore, MA, USA), and blot images were collected with a Bio-Rad ChemiDoc imaging system.

### RNA isolation and quantitative real-time RT-PCR analysis

Tissues/cells or organoids were lysed in RNA lysis buffer, and total RNA was extracted with the Purelink RNA mini kit (Invitrogen, Carlsbad, CA, USA) according to the manufacturer’s instructions. RNA was quantified with a Nano-Drop 2000 (Thermo Scientific, Waltham, MA, USA). Then, 1 μg of RNA was reverse transcribed with the High-Capacity cDNA Reverse Transcription Kit (Thermo Scientific, Waltham, MA, USA). The amplified cDNA was diluted to 10 ng/μl, and qRT-PCR was performed with primers listed in **Table S2** and TaqMan 2X master mix (Applied Biosystems/Life Technologies, Grand Island, NY) on a 7500 Fast Real-time PCR system (Applied Biosystems/Life Technologies). The delta-delta cycle threshold method was used to normalize expression to the reference gene 18S.

### RNA Sequencing Data Analysis

Sequencing quality and adapter contamination were assessed using FastQC v0.11.9 software. Adapter sequences were trimmed using fastp v0.20.0 software, and trimmed reads were aligned to the genome index using STAR v2.7.9a [STAR – see citation below]. The STAR genome index was created using raw FASTA and annotation files downloaded from the GENCODE portal for mouse genome build GRCm38 release V33. To account for unstranded reads, STAR – outSAMstrandField IntronMotif was enabled. Coordinate sorted binary format (BAM) files were generated for each output alignment file using SAMtools (46). Genome coverage tracks were created, using bamCoverage from the Deeptools software suite (47), with CPM normalization and a bin size of 5 bp. Summaries of read quality and alignment quality were generated using MultiQC v1.12 (48).

### Differentially Expressed Genes (DEGs)

Gene expression values were computed as the number of reads aligned per gene, using STAR—quantMode GeneCounts. The analysis for differential gene expression was carried out using DESeq2 (49). Raw counts were normalized, and genes with an average read count <10 across all samples were excluded from the differential analysis. A false discovery rate (FDR) cut-off of 0.05 and fold change cut-off of 20% (-0.263 ≤ log2(FC) ≥ +0.263) were used to identify differentially expressed genes (DEGs). Visualizations of DEGs were created in R v4.1.2 software using the and heatmap packages.

### Alternative Splicing Analysis

Alternative splicing events were determined using rMATS v4.1.2 (50), with STAR alignment files (BAM) and the GENCODE reference annotation (GTF) for mouse genome build GRCm38 release v33. Splicing events were classified into 5 categories: skipped exons, retained introns, mutually exclusive exons, alternative 5’ splice sites, and alternative 3’ splice sites. An FDR cut-off of 0.05 was used to screen for statistical significance, and events with an inclusion level difference (ILD) >= 0.2 or <= −0.2 were designated as including or excluding the exon of interest, respectively. The strongest events were then visually validated in integrative genomic viewer (IGV) (51) using bigwigs tracks created for each sample. Genome coverage tracks and sashimi plots were plotted using pyGenomeTracks (52).

### Gene Ontology

Over-representation analyses were performed in WebGestalt (WEB-based Gene Set Analysis Toolkit) (53) on lists of upregulated and downregulated DEGs and a list of genes with significant alternative splicing events. Significant enrichment was determined using an FDR cutoff of 0.05, with a Benjamini-Hochberg correction, and the minimum number of genes per category was set to 10. The top 6 strongest enrichment terms, by enrichment ratio, were extracted and visualized in R using ggplot2.

### PCR validation of MXE events

To validate the MXE AS events, RNA (typically 1000 ng) was reverse transcribed with the High-Capacity cDNA Reverse Transcription Kit (Thermo Scientific, Waltham, MA, USA). The amplified cDNA was diluted to 10 ng/μl, and PCR was performed using sequence specific primers listed in (see Supplemental Information for DNA sequence and annealing temperatures) **Table S2**. Routinely, 30 cycles of amplification were performed (94°C, 30 sec denaturation; 58°C, 30 sec annealing; 72°C, 60 sec extension) and samples were then analyzed by agarose gel electrophoresis. Images were recorded with a BioRad documenter.

### Quantification and statistical analysis

A two-tailed paired student t-test was used to analyze data from experiments with two experimental groups and one-way ANOVA followed by Tukey’s post hoc multiple range test was used for multiple comparisons. All data are presented as mean ± SE. GraphPad Prism 9 software was used for all statistical analyses. All the statistical details related to tests performed, sample size, and p-values can be found in the corresponding figure legends. To ensure reproducibility, technical and experimental rigor, and demonstrate biological significance of our findings, experiments were replicated in a minimum of three independent samples.

## Results

### SF3B1 is required for mouse embryo implantation and decidualization

Pioneering splicing factor SF3B1 is primarily implicated in human cancers, and we previously found an oncogenic role for this RNA-binding protein in endometrial cancer (9). However, the physiological role of this factor in any human physiologies is yet to be identified. Therefore, we sought to investigate whether SF3B1-dependent splicing functions in endometrial physiology. First, the expression of *Sf3b1* in the murine uterus during early pregnancy events was evaluated. Toward this, we mated female WT mice with males of known fertility and then stained their uteri with an SF3B1-specific antibody at different days in early pregnancy. In days 1–3, SF3B1 was predominantly expressed in uterine epithelia and glands (**Fig. 1A**). However, beginning on day 4, when the uterus is receptive to embryo implantation, strong SF3B1 staining was seen in the stroma. This staining was evident at least through day 7, which is when uterine decidualization occurs (**Fig. 1A**). However, we noted no difference in the abundance of *Sf3b1* mRNA at any stage in pregnancy (**Fig. 1B**). This analysis suggests a possible role for SF3B1 protein in the uterine decidualization program. To address this, we utilized PLAD-B, a macrocyclic splicing inhibitor that targets the SF3B1 subunit of the spliceosome (9, 54–56). Specifically, the binding of PLAD-B to SF3B1 stalls U2 snRNA in an open conformation to inhibit pre-mRNA splicing (57). Importantly, a point mutation in SF3B1 (R1074H) conferred resistance to PLAD-B cytotoxicity, indicating the stringent specificity of PLAD-B for SF3B1 (54). We intraperitoneally injected mice with vehicle or PLAD-B and then performed a commonly used artificial decidualization procedure in which one uterine horn is injected with sesame oil and the other horn is left intact (**Fig. S1A**). Oil-stimulated uterine horns from vehicle-treated mice were substantially larger (**Fig. 1C**) and had more proliferative, phospho-histone-H3-positive cells (**Fig. S1B**) than the control horns. In contrast, stimulated horns from PLAD-B-treated mice did not increase in size (**Fig. 1C**) and had fewer proliferative cells than horns from vehicle-treated mice (**Fig. S1B**). Also, the decidualization markers *Wnt4* and *Bmp2* increased significantly more in stimulated horns from vehicle-treated mice than in stimulated horns from PLAD-B-treated mice (**Fig. 1D**). Moreover, the stimulated horns from oil-treated mice had a greater percentage of SF3B1-positive stromal cells than the control horns did, which was the same in PLAD-B treated uteri (**Fig. S1C**). Since uterine stromal cell proliferation precedes decidualization during embryo implantation, we wondered whether SF3B1 is also required for embryo survival and implantation. To answer this question, we mated 6-8-week-old wild-type (WT) female mice with WT males of proven fertility. On the morning of day 3 of pregnancy, we intraperitoneally injected the mice with 5 mg/kg PLAD-B. PLAD-B-treated mice produced similar numbers of blastocysts on day 4 as vehicle-treated mice (**Fig. 1F**), indicating that PLAD-B did not impair embryo survival. However, the treatment dramatically impaired implantation, as PLAD-B-treated mice had significantly fewer implantation sites on day 5 of pregnancy than vehicle-treated mice (**Fig. 1E**). These data revealed a physiological role for SF3B1 in endometrial physiology, specifically embryo implantation and decidualization.

**Fig 1:**
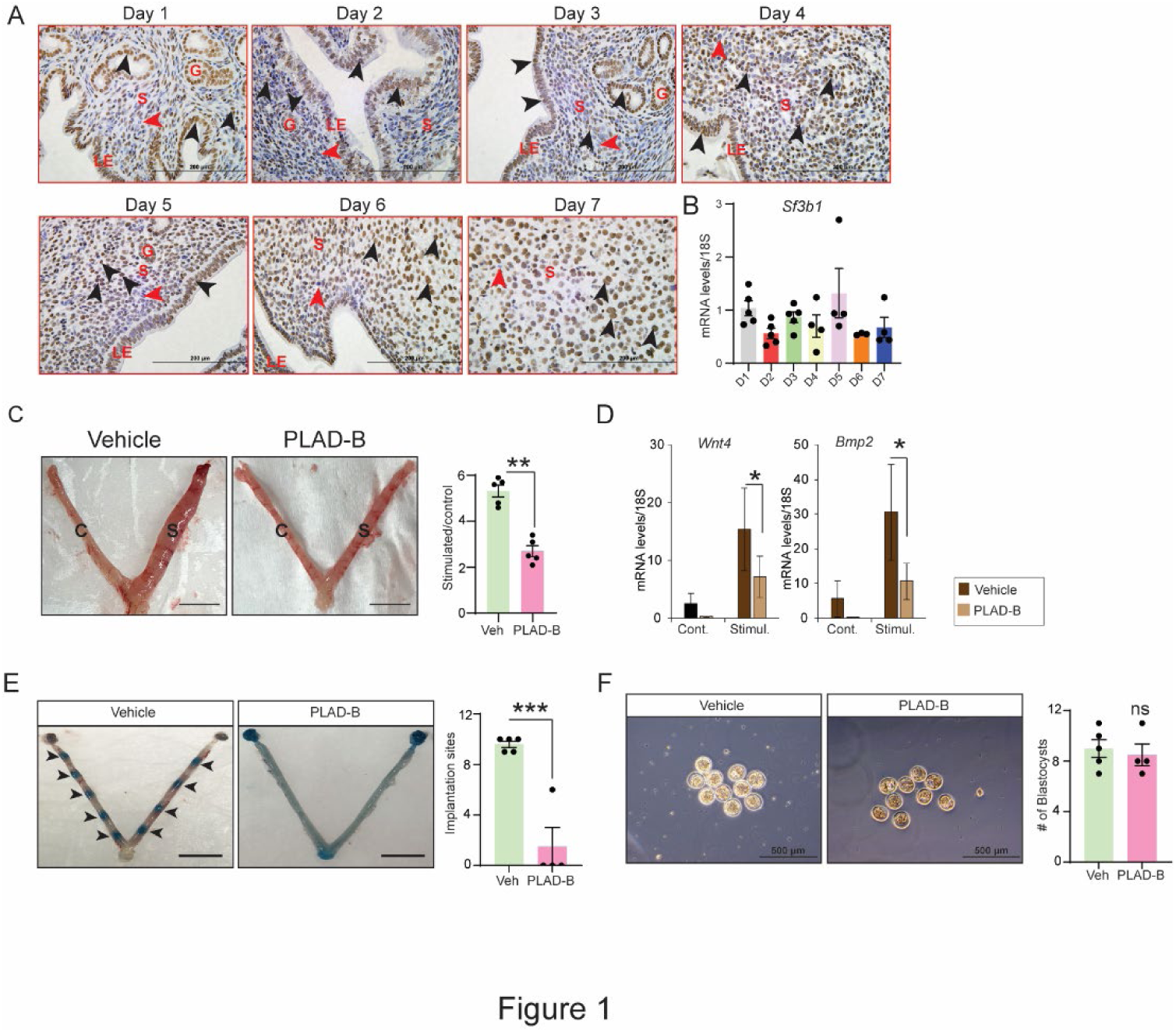
SF3B1 protein expressed in endometrium and is essential for uterine decidualization and embryo implantation in mice. **(A)** Representative images of uteri from mated mice at the indicated days of pregnancy stained with an SF3B1-specific antibody (brown). Black arrows, SF3B1-positive cells; red arrows, SF3B1-negative cells. LE: luminal epithelium, G: glands, S: stroma, Scale bar: 200 µm. (**B**) Relative abundance of *Sf3b1* mRNA in uteri from mated mice at indicated days of pregnancy. Data are presented as mean ±SE; n=5. (**C**) Gross morphology of uterus from hormone-treated vehicle or PLAD-B-treated mouse. Representative images and wet weights (graph) of control (C) and stimulated (S) uterine horns from vehicle-and PLAD-B treated mice. (**D**) Relative *Sf3b1*, *Wnt4, and Bmp2* transcript abundances in control and stimulated horns from vehicle-or PLAD-B-treated mice. Data are presented as mean ±SE (n=3). \**P*< 0.05, \*\**P*< 0.01. (**E**) Representative images of Chicago Sky Blue-stained uteri from vehicle-and PLAD-B-treated mice. Black arrowheads indicate implantation sites and graph depicting the numbers of implantation sites. (**F**) Representative images and numbers of blastocysts (graph on right) from vehicle-and PLAD-B-treated mice. Data are presented as mean ±SE. n=4. *P*> 0.05 as non-significant.

### SF3B1 is required for primary human decidualization in vitro

Given that SF3B1 is required for artificial decidualization in mice, we determined whether it is also required for HESC decidualization. Analysis of SF3B1 protein expression in proliferative and secretory phase endometrium from human subjects also indicated its robust expression in stromal cells (**Fig. 2A**). We further transfected HESCs with control or *SF3B1*-targeting siRNAs and treated with media containing estradiol, medroxyprogesterone acetate, and cyclic AMP (EPC, to induce decidualization). HESCs that received control siRNA changed from fibroblastic to epithelioid morphology (**Fig. 2B**) and had increased expression of the decidualization markers *IGFBP1* and *PRL* (**Fig. 2C**) by day three. In contrast, HESCs with *SF3B1* knockdown did not show a dramatic morphology change over six days (**Fig. 2B**) and expressed significantly less *SF3B1*, *PRL*, and *IGFBP1* than control cells (**Fig. 2C**). Consistent with in vivo findings, we noted no difference in the abundance of *Sf3b1* mRNA at any stage during in vitro decidualization process (**Fig. S2A**). Moreover, knockdown of *SF3B1* impairs primary HESC decidualization more than knockdown of other well-known splicing factors, *SF3B3, SRSF3* and *SF3A1* (**Fig. S2B**). To confirm this finding, primary HESCs treated with vehicle or EPC, or EPC plus 10 nM or 15 nM PLAD-B 24 hr prior to addition of EPC media. Whereas cells treated with EPC underwent a morphological change (**Fig. 2D**) and showed increased expression of the decidualization markers *PRL* and *IGFBP1* (**Fig. 2E**), these changes did not occur in cells treated with 10 or 15 nM PLAD-B. Furthermore, cells treated with PLAD-B prior to EPC failed to undergo decidualization (**Fig. S3A & B**), while cells treated with EPC after decidualization induction continued this cellular transformation process (**Fig. S3C & D**). These results indicate an indispensable role for SF3B1 in endometrial decidualization process.

**Fig 2:**
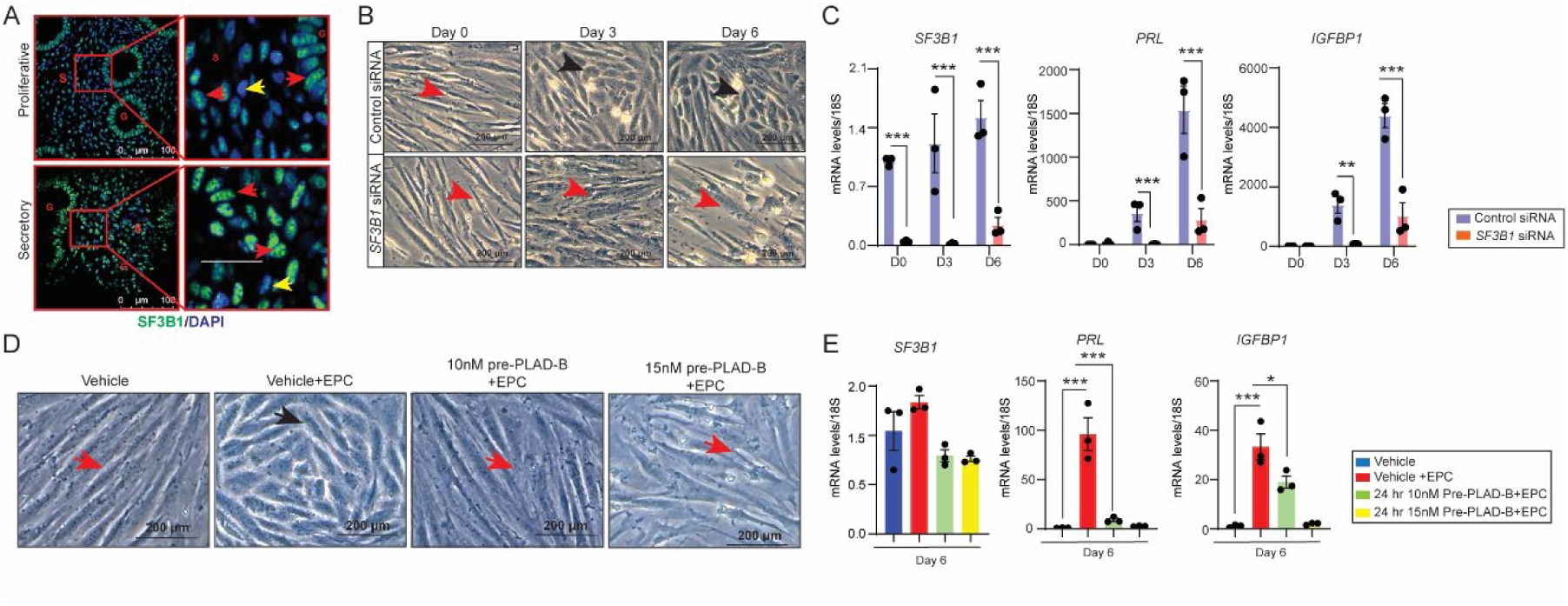
SF3B1 is essential for primary human endometrial stromal cell decidualization. (**A**) Immunofluorescence analysis to detect SF3B1 expression in endometrium from proliferative and secretory phase biopsies. Red arrows, SF3B1-positive cells; yellow arrows, SF3B1-negative cells. (**B**) Morphological changes and (**C**) transcript abundance of *SF3B1, PRL*, and *IGFBP1 from* HESCs transfected with control siRNA or *SF3B1* siRNA and cultured in decidualization media (EPC) for the indicated numbers of days. (**D**) Morphological changes and (**E**) transcript levels of *IGFBP1* and *PRL* of HESCs treated with vehicle or PLAD-B for 24 hrs and then treated with EPC cocktail for six days. Representative data from three technical replicates of one patient sample are shown as mean ±SEM; \*\**P* < 0.01 and \*\*\**P* < 0.001, scale bar, 200 µm.

### Identification of SF3B1-driven alternative splicing (AS) events in the endometrium

Given that the splicing function of SF3B1 is essential for endometrial decidualization, we next identified the mRNA splicing variants generated by SF3B1 in decidualizing HESCs. Three independent HESC lines (biological replicates) with three technical replicates of each HESCs cells line were transfected with control or *SF3B1* siRNA and treated with EPC. Knockdown of *SF3B1* was confirmed, total RNA was pooled from technical replicates and subjected to RNA-Seq to detect both RNA transcripts and splice variants in each group. *SF3B1* knockdown led to alterations in significant number of mRNAs (∼4500) in decidualizing HESCs (**Fig. 3A**). Specifically, a total of 3774 and 4410 mature mRNA went up or down regulated with *SF3B1* knockdown respectively, indicating a SF3B1 specific transcriptome in endometrium (**Fig. 3B**). Pathway analysis also revealed networks that promote RNA processing and cellular differentiation (**Fig. 3C**).

**Fig. 3:**
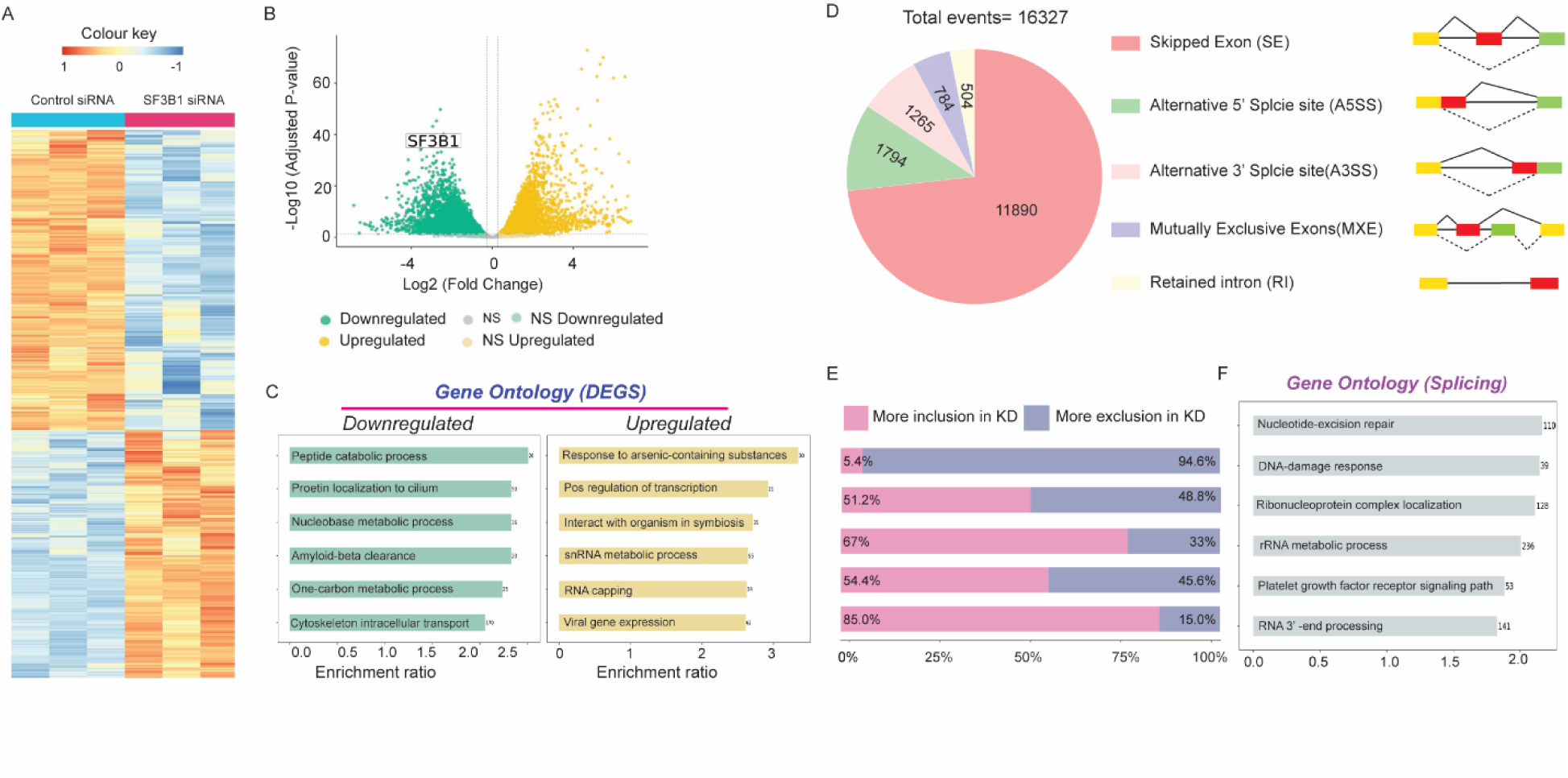
Identification of SF3B1-dependent differential transcriptome and alternative splicing during endometrial stromal cell decidualization. (**A-C**) SF3B1 transcriptome in decidualizing HESC. HESCs were transfected with *SF3B1* siRNA and cultured in decidualization media (EPC) for 72hrs and subjected to RNA-Seq. (**A**) Heat map of altered DEGs with SF3B1 knockdown changes (**B**) Volcano plot showing both the upregulated and downregulated gene distribution. (**C**) Gene ontology of enriched signaling pathways from SF3B1 transcriptome. (**D**) Pie chart depicting the total number of AS events altered with SF3B1 depletion. The AS events are classified into 5 categories: skipped exon (SE), retained intron (RI), alternative 5′ splice site (A5SS), alternative 3′ splice site (A3SS), and mutually exclusive exon (MXE). (**E**) The graph shows the % inclusion or exclusion in *SF3B1*-depleted cells for each alternative splicing event. (**F**) Gene ontology of enriched singling pathways from genes that are differentially altered with SF3B1-driven splicing.

Furthermore, we conducted analysis of alternative splicing analysis, adhering to a significance level of FDR <= 0.05 and an inclusion level difference ≥ 0.2 or ≤ −0.2. A total of 16327 alternative splicing (AS) events from 5808 genes were identified following SF3B1 knockdown in HESC cells (**Fig. 3D**). The SF3B1-dependent AS events encompassed five distinct types, with exon skipping being the most frequent, constituting 73% of all the AS events (**Fig. 3D**). Each of these five AS events were further distinguished into inclusion and exclusion events. Inclusion events refer to those with a positive Inc Level Difference, indicating higher exon inclusion levels in SF3B1 knockdown (KD) condition. Conversely, exclusion events are characterized by a negative Inc Level Difference, signifying lower exon inclusion levels in the *SF3B1* KD condition. Among the exon skipping events, a significant majority (95%) of the exons were excluded with *SF3B1* knockdown. Contrastingly, for the retained introns, we observed a distribution of 85% inclusion and 15% exclusion events. This balance between the exclusion of skipping exon (SE) events and the inclusion of retained intron (RI) events aligns with the known splicing signature of SF3B1(58). The alternative acceptor (A5SS) and alternative donor (A3SS) events exhibited an equal distribution of exon exclusion and inclusion. Intriguingly, we discovered that the loss of SF3B1 led to a significant number of mutually exclusive exon (MXE) AS events, a function not previously ascribed to SF3B1 (**Fig. 3D**). Functional enrichment analysis of these splicing (**Fig. 3E**) revealed DNA damage response and nucleotide-excision repair pathways (**Fig. 3F**). This comprehensive analysis unveils a unique SF3B1 dependent endometrium-specific alternative splicing landscape.

### SF3B1-driven MXE AS event is essential for stromal cell decidualization

Having identified a rare and previously unreported AS event (MXE) regulation by *SF3B1* in stromal cells, we next determined whether such MXE AS events are essential for stromal cell decidualization. Toward this, we chose four candidate genes with the highest degree of MXE AS events, *GLXR3*, *RBM39, ANAX5*, *CNOT9*, (**Fig. 4A & Fig. S4**). Using the read coverage, we identified the exons that are excluded or included in the respective MXE event in these genes (**Fig. 4A & D, S4A & D**). For example, in *RBM39* gene, knockdown of *SF3B1* resulted in inclusion of the second exon (chr20:35729461-35725037) with more exclusion of the first exon (chr20:35729311-35729365), indicating the MXE AS event occurrence (**Fig. 4D & E**). Next, using exon junctional analysis, we designed primers to validate these MXE events. Amplification of the differentially spliced exons with PCR clearly demonstrated that loss of *SF3B1* led to inclusion of one exon over the other (**Fig. 4C & F, S4C & F**). Since the SF3B1-driven MXE AS event regulates the mRNA variants, we next assessed whether these SF3B1-regulated genes also exert a biological action on *SF3B1*. To test this, we knocked down these four validated genes in HESCs and monitored the decidualization process (**Fig. 4G & S4G**). While knockdown of *RBM39*, *ANAX5* and *GLRX3* led to a significant block in fibroblastic-to-epithelioid morphological change, knockdown of *CNOT9* had a modest effect on this decidual cell transformation (**Fig. 4G & S4G**). Consistent with the morphological change, knockdown of these four genes led to significant downregulation in *PRL* and *IGFBP1* transcript levels (**Fig. 4H & S4H**). These data clearly suggest that SF3B1-driven MXE events contribute to decidualization process, possibly including the functional exons on effector genes such as RBM39.

**Fig. 4:**
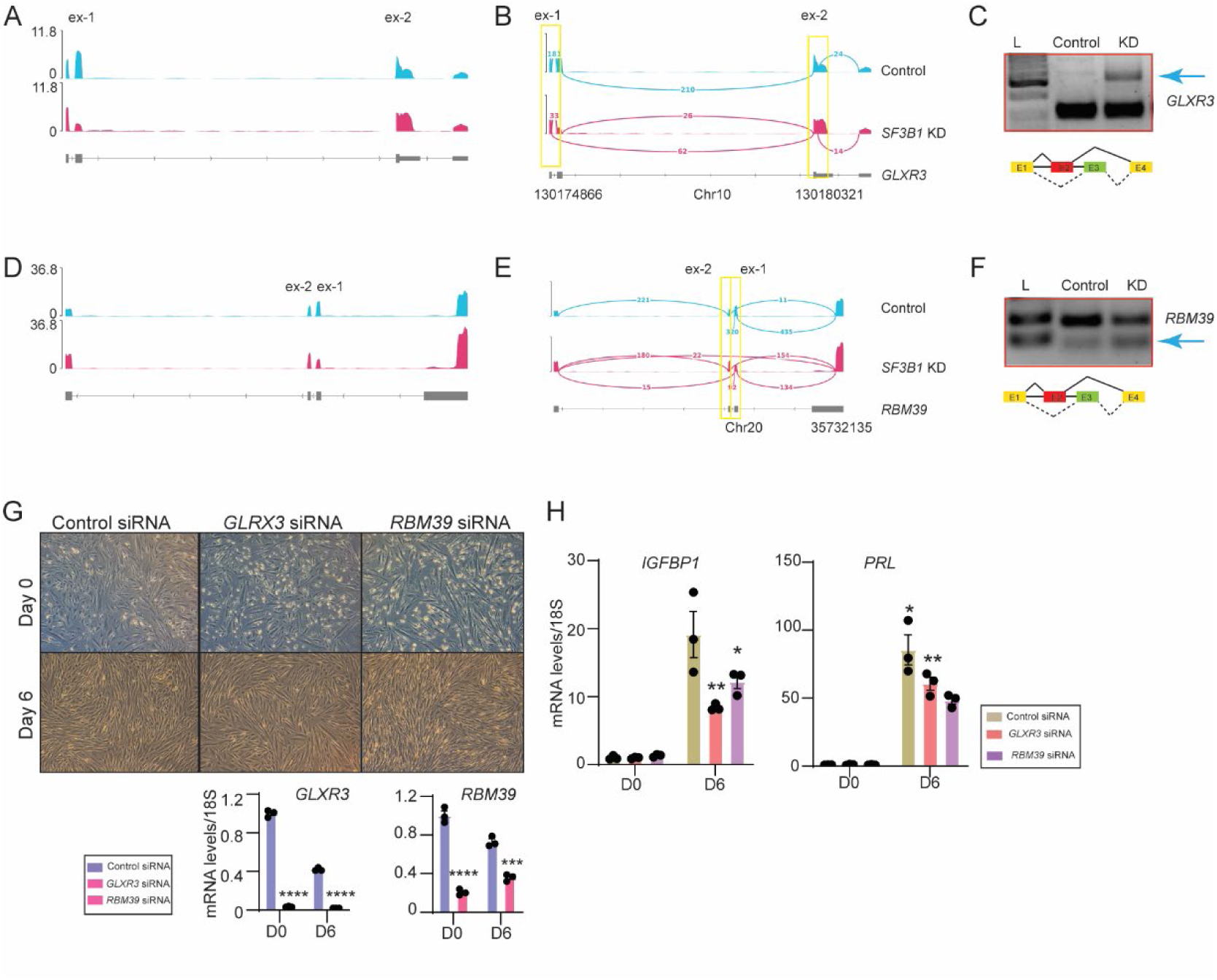
SF3B1-driven MXE AS events are essential for stromal cell decidualization. (**A & D**) Big wig and (**B & E**) sashimi plots depicting the MXE events in two candidate genes *GLXR3* and *RBM39* in which mutual exons are differentially used with *SF3B1* presence and absence. The constitutive splicing isoform is shown in the upper track in blue and the alternative splicing isoform is shown in the lower track in pink. (**C & F**) RT-PCR validation of *SF3B1*-affected MXE splicing events in *GLXR3* and *RBM39* candidate genes. Representative images from 3 independent experiments are presented. The illustration of each PCR product is indicated schematically right below the validation. (**G**) (*upper pane*l) Morphological changes and (*lower panel*) transcript abundance of *GLXR3, and RBM39 from* HESCs transfected with control siRNA or *GLXR3 or RBM39* siRNA and cultured in decidualization media (EPC) for the indicated numbers of days. (**H**) Transcript levels of *IGFBP1* and *PRL* in control siRNA or *GLXR3 or RBM39* siRNA transfected HESCs.

### Progesterone regulates SF3B1 protein stability in endometrial stromal cells via deubiquitinase enzyme

We consistently found that the SF3B1 protein was prominently detected in endometrial stromal cells during the human endometrium secretory phase (**Fig. 2A**). Since, progesterone (P4) drives the stromal cell decidualization, and its concentration is high in the secretory phase of the menstrual cycle, we investigated whether P4 regulates SF3B1 expression in stromal cells. First, we performed artificial decidualization in mice and found that the stimulated horns expressed more SF3B1 protein than control horns (**Fig. 5A & B**) but amounts of *Sf3b1* mRNA were similar between stimulated and control horns (**Fig. 5C**). To test whether in fact progesterone promoted this increase in SF3B1 protein, we treated primary HESCs with MPA and then measured SF3B1 protein and mRNA levels. Increased SF3B1 protein was evident as early as 6 hours after MPA treatment and remained elevated through at least 12 hrs (**Fig. 5D**). In contrast, *Sf3b1* transcript abundance was unchanged by MPA treatment (**Fig. 5E**). Finally, we investigated the mechanism(s) by which progesterone regulates SF3B1 in the endometrium. To determine the genomic and non-genomic functions of MPA on SF3B1 protein stabilization, we utilized a well-characterized antagonist of progesterone-PR genomic action, RU486. Treatment of HESCs with RU486 failed to counteract MPA-dependent SF3B1 protein stabilization, suggesting progesterone stabilizes SF3B1 protein possibly through non-genomic actions (**Fig. 5F**). Ubiquitin and ubiquitin-related proteins are central to the protein stability, thus we determined whether progesterone stabilizes SF3B1 through any deubiquitinase. Since, ubiquitin-specific peptidase 7 (USP7) deubiquitinated several splicing factors, the impact of USP7 inhibition on SF3B1 was measured. In primary HESCs, increased SF3B with MPA was completely abrogated with USP7 knockdown, suggesting that progesterone stabilizes SF3B1 through USP7 deubiquitinating signaling (**Fig. 5G**). Taken together, we found progesterone stabilizes SF3B1 and SF3B1-mediated mRNA splicing is crucial for progesterone-driven endometrial decidualization. These findings place SF3B1 in a feed-forward regulatory loop that mediates progesterone-induced transcriptional signals required for endometrial decidualization (**Fig. 5H**).

**Fig. 5:**
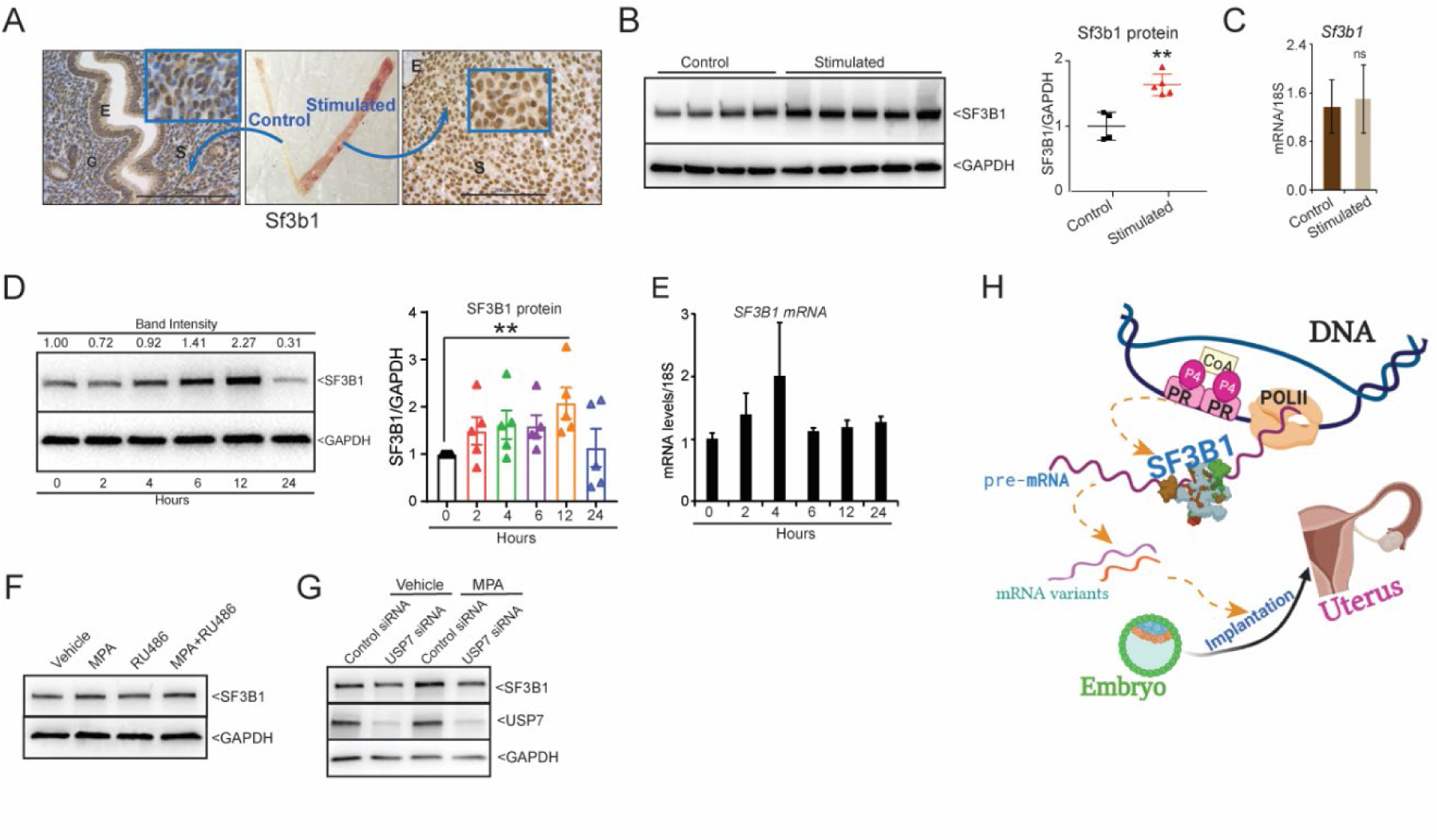
Progesterone regulates SF3B1 protein stability through USP7. (**A**) Representative uterus with control and stimulated horns (middle), and respective horns stained with an SF3B1. E, epithelium; G, gland; S, stroma. Scale bar: 200 μm. (**B**) Immunoblotting and quantification of SF3B1 protein and (**C**) qRT-PCR of *Sf3b1* mRNA in control and stimulated horns two days after the stimulus. n=4-5; \*\**P*<0.01. (**D-E**) HESCs treated with 1 µM MPA for the indicated times. (**D**) Representative immuno-blotting and quantification of SF3B1 protein from 5 different patient samples. (**E**) qRT-PCR of *SF3B1* mRNA abundance. Representative data from three replicates from one patient sample are shown as mean ±SEM. \*\**P*<0.01; n=5. (**F**) HESCs were treated with vehicle or RU486 for 2 hrs and then treated with MPA and immunoblotting of SF3B1 was performed. (**G**) HESCs transfected with control siRNA or *USP7* siRNA and treated with MPA for 12 hrs SF3B1 and USP7 protein levels measured through immunoblotting. (**H**) Schematic illustration of the hypothesis showing how P4/PR driven transcription might be coordinating with SF3B1-dependent splicing to generate different mRNA variants which are essential for successful embryo implantation in uterus.

## Discussion

To establish pregnancy, the endometrium undergoes intricate cellular transformation events including cell proliferation, differentiation, and apoptosis. These changes are coordinated by cell type-specific responses to the steroid hormones, estrogen and progesterone. Although much of the transcriptional repertoire driven by these two hormones in the endometrium is well-established, we lack a complete picture of the downstream processing of these mRNA transcripts, including the mRNA splicing. In this study, we found splicing factor SF3B1 mediates progesterone-driven alternative splicing that is essential for uterine receptivity and decidualization. Notably, we show that abrogation of SF3B1 activity via PLAD-B treatment impaired stromal cell decidualization in both mice as well as human endometrial cells. We also show that SF3B1-dependent splicing paradigm is critical for decidualization. Finally, we found that progesterone acts as an upstream regulator and maintains high SF3B1 levels in mice as well as human endometrial cells possibly via deubiquitinating enzymes.

Splicing factor dysregulation can lead to the production of aberrant mRNA variants that contribute to cellular dysfunction in several pathologies (10–22). Evidence from mice and drosophila models did find that alternative splicing is an important developmental regulatory mechanism and is frequent during early embryonic development (59–62). Similarly, alternative splicing was reported to play a crucial role in tissue morphogenesis of the heart, and brain (63, 64). However, there is a significant knowledge gap on the tissue or cell-type specific biological actions of splicing factors. Our findings on an indispensable role of SF3B1 in stromal cell differentiation highlight a novel tissue-specific role for SF3B1 throughout biology. During the decidualization phase, stromal cells undergo cell cycle arrest and show proliferation inhibition by marking the onset of differentiation. Recently, the potential link between SF3B1 and cell-cycle progression gained attention when it was shown that phosphorylation of SF3B1 is cell cycle-dependent (65, 66). Importantly, it was demonstrated that nucleosome binding of SF3B1 is dynamic owing to its phosphorylation status, and thus SF3B1 regulates the alternative splicing of cell-cycle genes. For example, SF3B1 phosphorylation peaks at G2/M in a CDK1-dependent manner, thereby decreasing its binding with nucleosomes. In contrast, the interaction of SF3B1 with nucleosomes increases during G1/S when it has the lowest level of phosphorylation, owing to PP2A and PP1 phosphatase activities as the cell progresses through G1/S phase. A previous report by Logan et al., showed that endometrial stromal cells undergo cell cycle arrest at G0/G1 phase initially and G2/M phase at later stages during the time of decidualization (67). Moreover, impaired decidualization in our study when SF3B1 activity is abrogated prior to initiation or during the decidualization phase in mice and human endometrial stromal cells suggests that SF3B1 might be strictly involved in coordinating with cell cycle regulators to govern the splicing of specific target genes essential for decidualization. Interestingly, we found that progesterone (P4) promotes SF3B1 protein stability but not the mRNA levels. This is consistent with the fact that stabilization of SF3B1 by P4 appears through non-genomic actions. Thus, stabilization of SF3B1 by P4 non-genomic action might prime the cells for mRNA splicing. Specifically, the P4/PR-driven transcription might be coordinated with SF3B1-dependent splicing to generate vital mRNA variants. Furthermore, activated SF3B1 might act independently of PR to generate mature mRNA, which is essential for mediating P4 responses. However, substantial biochemical studies are needed to understand the transcriptional co-coupling between PR and SF3B1 in the endometrium.

We found that loss of SF3B1 function in stromal cells resulted in aberrations in alternative splicing events, resulting in the development of numerous aberrant transcripts curtailing the decidualization program. Although exon skipping and intron retention are the two key AS events driven by SF3B1 (11, 68–72), we found SF3B1 is regulating MXE AS events in stromal cells. Given, majority of the AS event distributions for SF3B1 are reported from pathologies, regulation of MXE AS event thus underscores the tissue and/or cell-type specific actions of SF3B1 splicing factor. Consistently, we found SF3B1-driven MXE events in candidate mRNA, including *GLXR3*, *RBM39*, *ANXA5*, and *CNOT9* does not fully phenocopy SF3B1 function in stromal cells. These findings suggest that a functional impact of a single splicing factor can’t be attributed to a single AS event or a single mRNA variant. Instead, SF3B1 controls broad AS events to generate mature mRNA from pre-MRNA and mRNA variants to exert its biological functions, such as decidualization. Collectively, our study demonstrates that SF3B1-mediated alternative splicing is critical for decidualization and pregnancy establishment. Thus, uncovering the transcript variants that are associated with early pregnancy loss can be used to develop new strategies to diagnose or prevent early pregnancy loss.

### Non-standard Abbreviations

WT: Wild Type
MPA: Medroxy Progesterone Acetate
HESCs: Human Endometrial Stromal Cells
Dpc: Days Post Coitum
E2: Estrogen
P4: Progesterone
AS: Alternative Splicing
MXE: Mutually Exclusive Exons
ES: Exon Skipping

## Acknowledgments

We thank the Genome Technology Access Center in the Department of Genetics at Washington University School of Medicine for service with transcriptome analysis. The Center is partially supported by National Cancer Institute (NCI) Cancer Center Support Grant # P30CA91842 to the Siteman Cancer Center and by Institute for Clinical and Translational Science (ICTS)/ Clinical and Translational Sciences Award (CTSA) Grant # UL1TR002345 from the National Center for Research Resources, a component of the National Institutes of Health (NIH), and NIH Roadmap for Medical Research. We thank Dr. Robert Lawrence for editorial assistance with the manuscript.

## Author contributions

P.P., S.B.C., and R.K. designed experiments, conducted most of the studies, interpreted the data, and wrote the manuscript. M.D., and H.Y. helped with splicing data analysis. S.J.G., and P.J. provided critical reagents for the study. R.K. conceived the project, supervised the work, and wrote the manuscript. All authors critically reviewed the manuscript.

