## Supplementary Information for "SF3B1-dependent alternative splicing is critical for maintaining endometrial homeostasis and the establishment of pregnancy"

Alkek Center for Metagenomics and Microbiome Research

Baylor College of Medicine

Houston, TX, 77030

**Short title**: SF3B1 function in endometrial functions.

[*] These authors contributed equally

The authors have declared that no conflict of interest exists.

**Supplementary Table S1:** List of antibodies used.

| Antibody | Company, Catalogue number | Application |
|---|---|---|
| SF3B1 | Abcam, ab170854 | Immunoblotting, Immunohistochemistry Immunofluorescence |
| USP7 | Proteintech | Immunoblotting |
| Phospho-Histone H3 | Millipore Sigma, 06-570 | Immunohistochemistry |
| Normal Rabbit IgG | CST, #2729 | Immunohistochemistry |
| Goat anti-Rabbit IgG (H+L) | Thermofisher Scientific, A32731 | Immunofluorescence |
| GAPDH | CST, #2118S | Immunoblotting |
| Anti-rabbit IgG, HRP-linked | CST, #7074 | Immunoblotting |

**Supplementary Table S2:** List of primers and TaqMan probes used.

| Gene name | Species | Application, Chemistry | Company | Sequence/Cat. No. |
|---|---|---|---|---|
| *Sf3b1* | Mouse | qPCR, Taqman | ABI | Mm01255457_m1 |
| *Sf3b1* | Mouse | qPCR, Taqman | ABI | Hs00961640_g1 |
| *IGFBP1* | Human | qPCR, Taqman | ABI | Hs00236877_m1 |
| *PRL* | Human | qPCR, Taqman | ABI | Hs00168730_m1 |
| *Lif* | Mouse | qPCR, Taqman | ABI | Mm00434762_g1 |
| *Wnt4* | Mouse | qPCR, Taqman | ABI | Mm01194003_m1 |
| *Bmp2* | Mouse | qPCR, Taqman | ABI | Mm01340178_m1 |
| *RBM39* | Human | qPCR, Taqman | ABI | Hs00863502_g1 |
| *GLRX3* | Human | qPCR, Taqman | ABI | Hs01582641_g1 |
| *RBM39* | Human | qPCR, Taqman | ABI | Hs01103365_m1 |
| *ANXA5* | Human | qPCR, Taqman | ABI | Hs00996186_m1 |
| *18S* | Mouse | qPCR, Taqman | ABI | 4318839 |
| RBM39 Exn-2 Forward | Human | PCR | Sigma | CTTAATTTGATGCTATGAGGCAACC |
| RBM39 Exn-2 Reverse | Human | PCR | Sigma | TAACAGTGCCATCCGAGGAA |
| RBM39 Exn-1+2 Forward | Human | PCR | Sigma | CTCACAGGGCTCTTGTCTTT |
| RBM39 Exn-1+2 Reverse | Human | PCR | Sigma | TCCGAGGAAAGATTGGGTTG |
| GLRX3 Exn-1 Forward | Human | PCR | Sigma | ATACCCTCAGCTGTATGTGAAAG |
| GLRX3 Exn-1 Reverse | Human | PCR | Sigma | CAATATCCAATCCTCCCACCAG |
| GLRX3 Exn-1+2 Forward | Human | PCR | Sigma | ACCCTCAGCTGTATGTGAAAG |
| GLRX3 Exn-1+2 Reverse | Human | PCR | Sigma | TAGTTGGGCACCAAGTTTAAGA |
| GLRX3 Exn-2 Forward | Human | PCR | Sigma | ATGGTGAATTGCTGCCTATACT |
| GLRX3 Exn-2 Reverse | Human | PCR | Sigma | TCATGAACTGCTCCCAATGTAA |
| CNOT9 Exn-1 Forward | Human | PCR | Sigma | CGTTTCTCGCAGCACACAT |
| CNOT9 Exn-1 Reverse | Human | PCR | Sigma | GAGATACTCAAAGGGACGTGTTT |
| CNOT9 Exn-1+2 Forward | Human | PCR | Sigma | TGCACACTGTCAGCAAA |

| | | | | |
|---|---|---|---|---|
| CNOT9 Exn-1+2 Reverse | Human | PCR | Sigma | AGTTCACTTCCAGATTCCATAA |
| CNOT9 Exn-2 Forward | Human | PCR | Sigma | CAGAAATTATCCCTTTATGTTTGCG |
| CNOT9 Exn-2 Reverse | Human | PCR | Sigma | GAAAGTTCACTTCCAGATTCCATAA |
| ANXA5 Exn-1 Forward | Human | PCR | Sigma | CCTTCATAGCATAATAGAGGGTCTC |
| ANXA5 Exn-1 Reverse | Human | PCR | Sigma | TCGAAGTATACCTGCCTACCT |
| ANXA5 Exn-1+2 Forward | Human | PCR | Sigma | CCACTTACCTTCATAGCATAATAGAG |
| ANXA5 Exn-1+2 Reverse | Human | PCR | Sigma | GGATTTCAAATTGAGGAAACCATTG |
| ANXA5 Exn-2 Forward | Human | PCR | Sigma | CTACCAACAGCAAGGAGTAGTT |
| ANXA5 Exn-2 Reverse | Human | PCR | Sigma | TCAAATTGAGGAAACCATTGACC |

*all primer sequences are written 5’ to 3’
ABI-applied biosystems

**Fig. S1: SF3B1 is essential for stromal cell proliferation and transformation during uterine decidualization.** (**A**) The experimental strategy followed for hormone induction in the ovariectomized mouse. (**B**) Immunohistochemical detection of phospho-histone H3 (pH3) and (**C**) Sf3b1 in the stimulated horns tissues of the vehicle or PLAD-B treated mice. Black arrowheads indicate pH3-and Sf3b1 positively stained ESCs, and the red arrows show pH3/Sf3b1-negative cells.

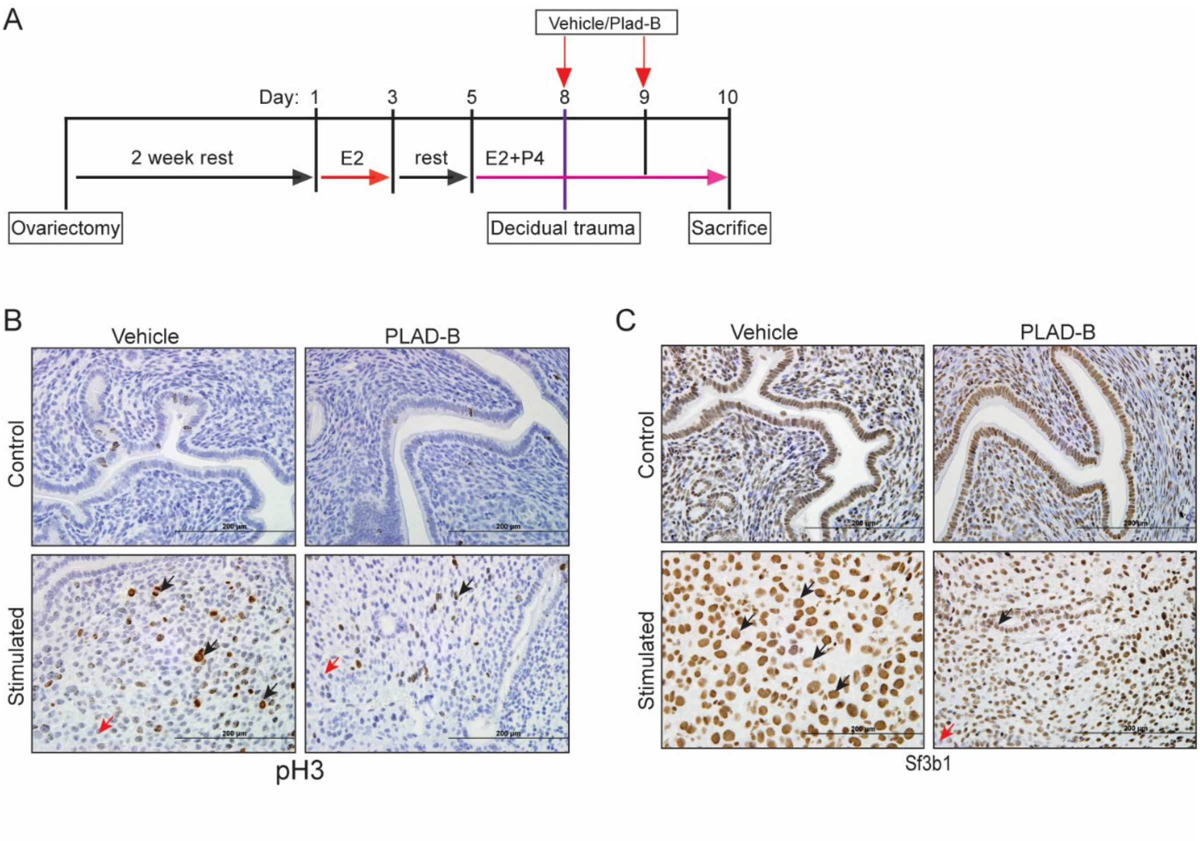


Fig. S1

**Fig. S2: SF3B1 is required for primary human endometrial stromal cell decidualization.** (**A**) Relative transcript levels of *SF3B1*, *IGFBP1,* and *PRL* in HESCs treated with EPC media for the indicated number of days. (**B**) HESCs were transfected with control siRNA or the indicated siRNA and cultured in decidualization media (EPC) for the indicated number of days. Abundances of PRL and IGFBP1 transcripts were measured. Representative data from three technical replicates of one patient sample are shown as mean ±SEM; *P < 0.05 and ***P < 0.001; n=3.

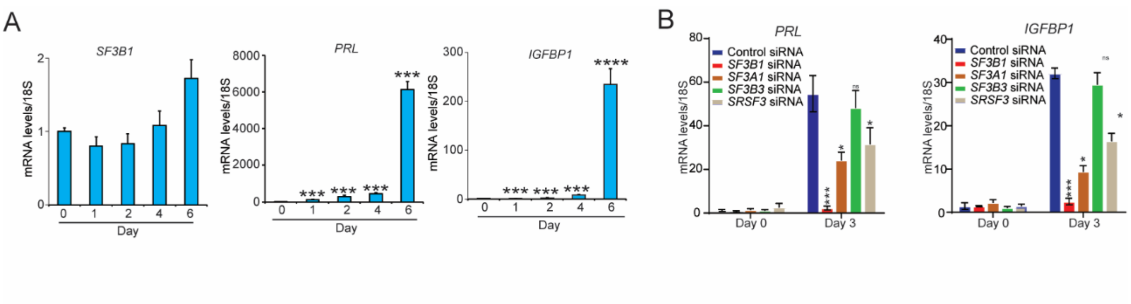


Figure S2

**Fig. S3: PLAD-B mediated inhibition of SF3B1 activity abrogates primary human endometrial stromal cell decidualization.** (**A**) Morphological changes in HESCs treated with vehicle or EPC cocktail with PLAD-B for six days of decidualization or (**C**) EPC media with PLAD-B starting from Day 3 to Day 6 of decidualization. (**B & D**) Relative transcript abundance of *SF3B1*, *PRL*, and *IGFBP1* in HESCs treated with EPC and PLAD-B at the indicated number of days. Representative data from three technical replicates of one patient sample are shown as mean ±SEM; $^{**}P < 0.01$ and $^{***}P < 0.001$, scale bar, 200 μm.

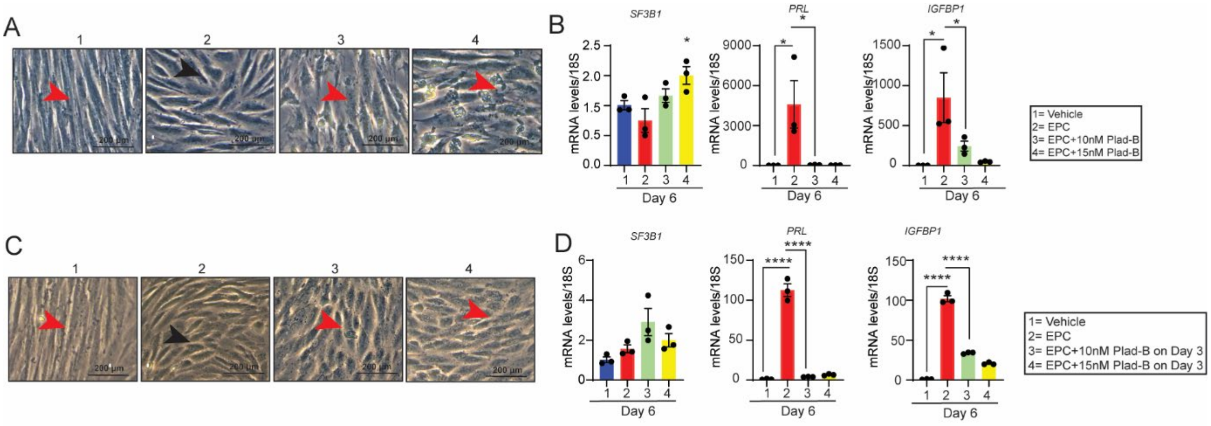


Figure S3

**Fig. S4: SF3B1-mediated MXE AS events are important for stromal cell decidualization.** (**A & D**) Big wig and (**B & E**) sashimi plots depicting the MXE events in two candidate genes *ANXA5* and *CNOT9* with differential mutual exon usage in the presence or absence *of SF3B1*. The constitutive splicing isoform is shown in the upper track in blue and the alternative splicing isoform is shown in the lower track in pink. (**C & F**) RT-PCR validation of SF3B1-affected MXE splicing events in *ANXA5,* and *CNOT9* candidate genes. Representative images from 3 independent experiments are presented. The illustration of each PCR product is indicated schematically right below the validation. (**G**) (*upper pane*l) Morphological changes and (***lower panel***) transcript abundance of *ANXA5, and CNOT9 from* HESCs transfected with control siRNA or *ANXA5 or CNOT9* siRNA and cultured in decidualization media (EPC) for the indicated numbers of days. (**H**) Transcript levels of *IGFBP1* and *PRL* in control siRNA or *ANXA5 or CNOT9* siRNA transfected HESCs.

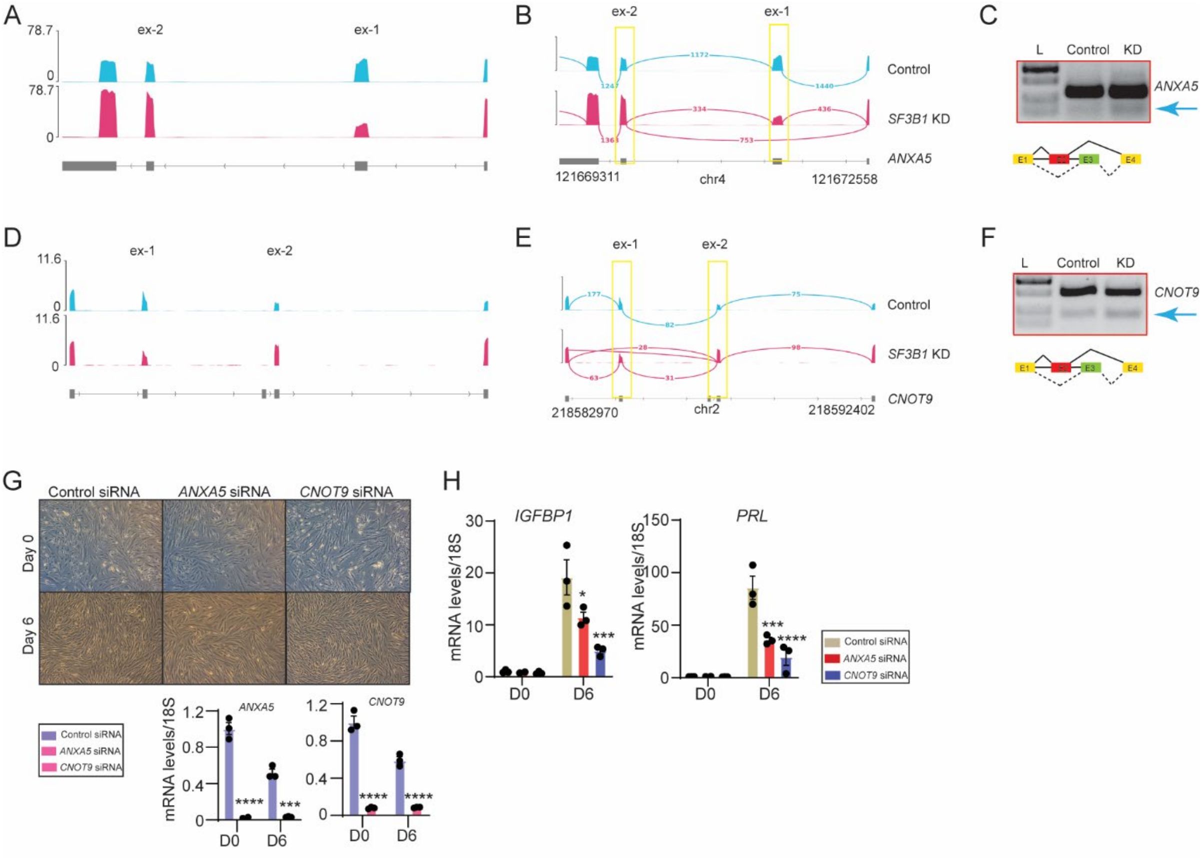


Figure S4
